# CellMap: Characterizing the types and composition of iPSC-derived cells from RNA-seq data

**DOI:** 10.1101/2021.05.24.445360

**Authors:** Zhengyu Ouyang, Nathanael Bourgeois, Eugenia Lyashenko, Paige Cundiff, Patrick F. Cullen, Ravi Challa, Kejie Li, Xinmin Zhang, Fergal Casey, Sandi Engle, Baohong Zhang, Maria I. Zavodszky

## Abstract

Induced pluripotent stem cell (iPSC) derived cell types are increasingly employed as *in vitro* model systems for drug discovery. For these studies to be meaningful, it is important to understand the reproducibility of the iPSC-derived cultures and their similarity to equivalent endogenous cell types. Single-cell and single-nucleus RNA sequencing (RNA-seq) are useful to gain such understanding, but they are expensive and time consuming, while bulk RNA-seq data can be generated quicker and at lower cost. *In silico* cell type decomposition is an efficient, inexpensive, and convenient alternative that can leverage bulk RNA-seq to derive more fine-grained information about these cultures. We developed CellMap, a computational tool that derives cell type profiles from publicly available single-cell and single-nucleus datasets to infer cell types in bulk RNA-seq data from iPSC-derived cell lines.

## Introduction

Human induced pluripotent stem cells (iPSC) and cells developed from them are gaining widespread acceptance for their use in understanding normal and disease processes *in vitro*^*1, 2*^. Derived directly from patient material with an unlimited capacity to proliferate in the undifferentiated state and the theoretical ability to differentiate into any cell type under appropriate experimental conditions, iPSCs and their differentiated derivatives represent an attractive alternative to traditional *in vitro* models that rely on cancer cell lines or rodent material. Barriers that once limited the use of iPSCs, such as costly reagents, complicated culture protocols, or restricted access to high quality, well-characterized iPSC lines, have diminished over the last decade. It has also become increasingly apparent that human biology can diverge significantly from rodent and even non-human primate biology, thus necessitating the use of human cells^3, 4^.

For *in vitro* studies using iPSC-derived cells to be meaningful, it is important to understand the relationship of the *in vitro* generated cells to endogenous cell types. This is particularly true during the development of novel iPSC differentiation protocols when seemingly small changes in culture conditions can lead to divergent cell fates. Evaluating expression of only a few canonical proteins by immunocytochemistry or genes by qPCR may not provide an adequate representation of what is in the cell culture dish and it can be technically challenging by these methods to test for the presence of off-target cell fates. A similar issue arises once a differentiation protocol has been established and one would like to ensure the reproducibility of specific cell type production within the same iPSC line and across different iPSC lines differentiated with the same protocol. While iPSC-derived cell types need not be perfect replicas of the endogenous cell type to be useful for disease modeling, they must reproduce the mechanism of the endogenous function to be evaluated in an appropriate cellular context to be physiologically relevant.

A variety of cell types exist in biological tissues performing different functions. When a biological function is altered or deficient, we need to understand the origin and mechanism of this aberration to devise ways of correcting it. However, most of the readily available biological samples are composed of a mixture of cell types. Experimentally separating these cell types and performing single-cell sequencing on them is cumbersome and costly. Therefore, developing computational approaches for cell type deconvolution from bulk RNA-seq data has been a popular and fruitful endeavor during the past decade^5, 6^. The earlier methods relied on cell type marker genes or cell-specific signatures obtained from prior publications or derived from low-throughput experiments, such as gene expression profiling on FACS sorted cells. CellMix was one such useful toolset that provided access to multiple deconvolution methods allowing the user to select the best approach based on the available data^7^. Unfortunately, the support of this public tool has been discontinued and it is incompatible with the latest R libraries. More complex and computationally demanding approaches have also been designed to characterize engineered cells based on inference of tissue-specific gene regulatory networks first from microarray data and then from bulk RNA sequencing data^8, 9^.

The aspiration to eliminate the dependence on prior knowledge of cell type markers or expression signatures led to efforts to develop *de novo* deconvolution algorithms. One such example is CellDistinguisher, which mathematically identifies cell type specific patterns in bulk expression data obtained from multiple heterogeneous samples^10^. CellDistinguisher can identify the genes that best distinguish a defined number of cell types or biological processes in the input data. This type of unsupervised deconvolution, however, cannot tell whether the patterns detected are a result of the presence of multiple cell types or subpopulations undergoing different biological processes, e.g., apoptosis, different phases of the cell cycle, etc. Adding even limited amount of prior knowledge in the form of marker gene sets can guide such methods into the desired direction.

With the advances of single-cell and single-nucleus (sc/sn) RNA sequencing, generating the much-needed prior knowledge to characterize cell types is gradually becoming a reality^11, 12^. These sequencing approaches are still quite expensive and arduous. Consequently, most laboratories cannot afford to apply them regularly to characterize iPSCs. But, given even a limited number of such datasets for the cell types of interest, they can be leveraged to mathematically characterize many samples for which the much easier and more cost-effective bulk RNA-seq data can be generated.

There are multiple deconvolution methods that rely on single-cell information to generate cell type expression profiles. Providing a thorough review of the field is beyond the scope of this work. Here we focus on a few representative examples that we used to benchmark our method. MuSiC uses scRNA-seq data to estimate cell type composition in samples with bulk RNA-seq data^13^. One limitation of this approach is that the query bulk needs to have the same cell types as the samples from which scRNA-seq data is derived. A more practical method would allow cell type profiles to be derived from sc/sn RNA-seq datasets of samples with disparate compositions of cell types. Bisque is designed to perform a very fast decomposition using non-negative least squares (NNLS) regression based on one single-cell dataset serving as a reference^14^. The constraint with this method, like with MuSiC, is that the user should have a single-cell dataset with cell types matching the bulk samples available before applying it. The most recent deconvolution method, SCDC derives expression profiles from multiple scRNA-seq datasets adopting an ensemble framework to implicitly address the batch effects inherent in datasets coming from different sources^15^. It achieves this by applying different weights to different datasets. The reference data most similar to the bulk data overall will have a higher weight. A drawback of this method is that all reference datasets need to have the same cell types. Meeting this condition is unlikely when using publicly available datasets from multiple sources.

For accurate disease modeling using human iPSCs, we needed a deconvolution tool designed to overcome the limitations encountered in existing tools. The new tool should be able to characterize a variety of cell types while making it possible to expand the list as needed. It should easily incorporate new reference datasets as they become available, allowing the user to retrain and retest the tool without modifying the code. To mitigate the batch effects or other biases inherent in individual datasets, a requirement has been imposed to have each cell type represented in at least three datasets, but we do not impose a constraint that each reference dataset need contain all cell types of interest.

CellMap was developed to meet these requirements. It was aimed mainly at characterizing iPSC-derived cells in terms of their cell type composition, their similarity to previously characterized primary cells or other iPSC-derived datasets, as well as assessing batch-to-batch variability. One important feature of this tool is the ability to regenerate the cell type profiles easily as newer or better-quality single-cell and single-nucleus datasets are produced and published. Here we demonstrate that besides iPSC-derived cells, CellMap can naturally be applied to the deconvolution of any complex samples whose constituent cell types are represented in the provided reference datasets.

## Methods

### Workflow

CellMap employs NNLS regression to decompose a bulk sample into cell type proportions based on the gene expression values (G) of a query bulk sample and the cell type specific expression profiles as shown in Eq (1):

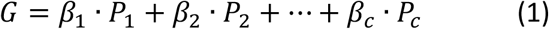

Where *G* is a *g* × 1 query expression vector with g denoting the number of genes. *P*_*1*_, *P*_*2*_, *…, P*_*c*_ are expression profiles (also *g* × 1 vectors) of *c* pure cell types derived from single cell or single-nucleus RNA-seq datasets, while *β*_*1*_, *β* _*2*_, *…, β* _*c*_ are the derived non-negative proportions of these cell types in the bulk sample. The overall workflow consists of two major steps: generating the cell type profiles and deconvolution of the query bulk sample (Fig. 1). While deriving the cell type profiles is the more computationally expensive part, it only needs to be carried out once for a given set of input sc/sn datasets.

**Figure 1.**
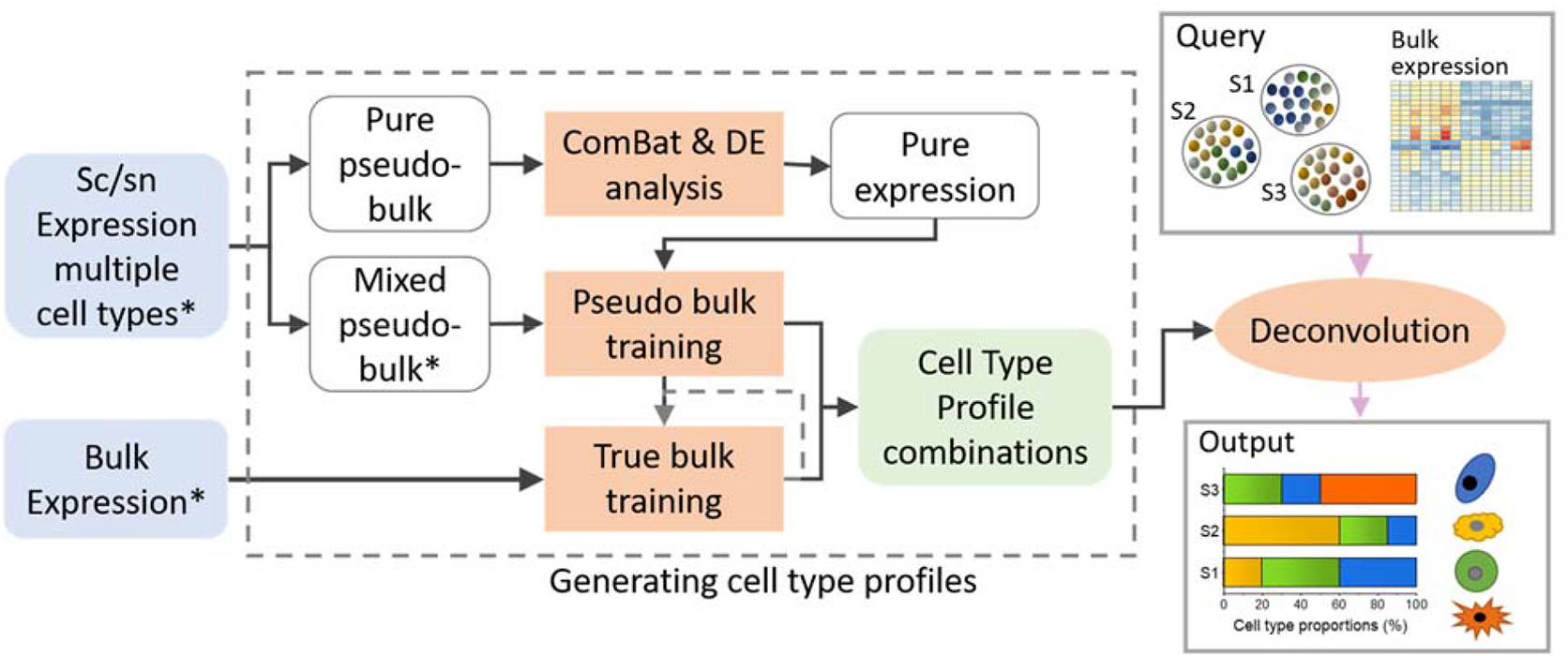
Overview of the CellMap workflow. CellMap includes the steps to generate the cell type specific profiles and the deconvolution which uses these profiles to predict the composition of a bulk sample. Datasets with samples of known cell-type compositions are denoted by *.

### Generating pseudo-bulk samples

The cell type expression profiles were derived from publicly available sc/sn RNA-seq datasets with cell type annotations provided by the authors (Supplementary Table 1). For skeletal muscle, we could only find bulk RNA-seq data and used those as is they were pseudo-bulk generated from single cells to derive the profiles. Since no two cells are identical, even if they are labeled as the same type, we attempted to capture this inherent biological heterogeneity by generating multiple pure cell type pseudo-bulk samples from each dataset. Cells were randomly selected from the existing pool of a given type and their expression values were summed up across the cells for each expressed gene. Similarly, mixed cell type pseudo-bulk samples with known cell type compositions were also generated from each dataset for the profile selection training. After the pseudo-bulk samples were normalized to 1M reads, genes with low expression were filtered out. Low expression genes were defined as having less than 4 counts in more than 20% of pseudo-bulk samples of any cell type. The expression metric used for full length transcript datasets was TPM (Transcripts Per Million), while UMI (Unique Molecular Identifier) was used in the case of datasets obtained with 3’ RNA sequencing.

### Normalization across datasets

ComBat normalization was used to eliminate batch effects among pure pseudo-bulk samples across reference datasets that can occur due to the differences in sequencing library preparation and sc/sn sequencing platforms^16^. Depending on the extent of overlapping cell types among datasets, two different strategies were applied. If shared cell types among datasets were sparse (Fig. 2A), ComBat was applied to the pseudo-bulk pure samples independently for each cell type across their source datasets. However, in the preferred scenario, when the overlap of cell types among the datasets was high (Fig. 2B, C), the normalization was performed across all pseudo-bulk samples generated for the pure cell types in one step. This approach was taken to avoid removing true cell type differences while performing batch correction for unbalanced datasets.

**Figure 2.**
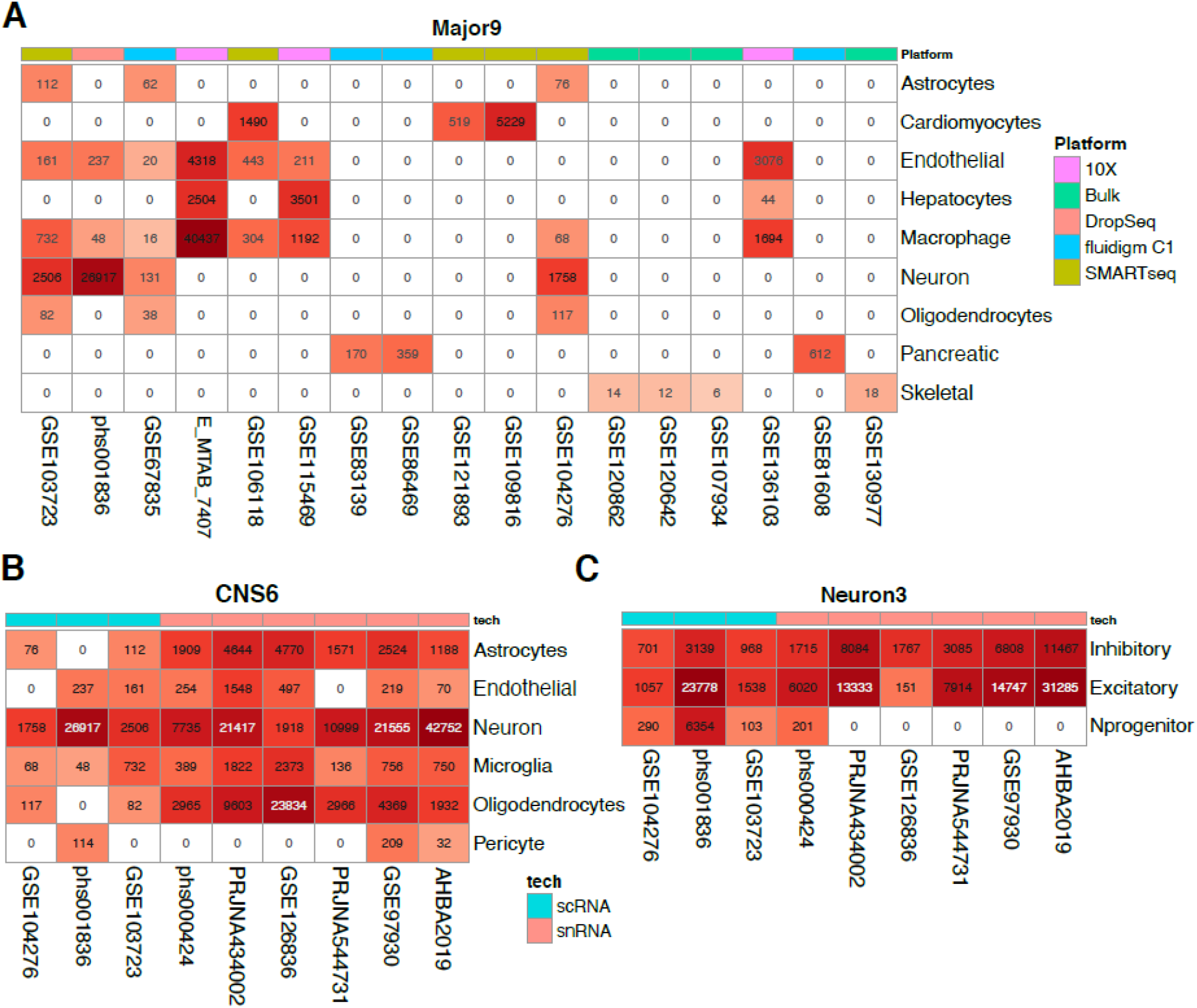
Single-cell and single-nuclei datasets used for cell type profile generation. (A) Datasets used for the 9 major cell types. (B) Datasets used for the 6 CNS cell types. (C) Datasets used for the neuronal subtypes of interest. The numbers in the colored boxes indicate the number of cells contained in each category after curation.

### Cell type profile genes

To increase the sensitivity of cell type detection, only cell type specific genes were included in the cell type profiles. Such genes were more highly expressed in one cell type relative to all others and they were identified by performing pairwise differential expression analysis with edgeR on the batch-corrected pseudo-bulk data^17^. For a balanced approach, an attempt was made to maintain a similar number of profile genes for all cell types within a group (see description of cell type groups under Stepwise Deconvolution below). Thus, fewer genes were kept for the CNS6 profiles than those of Major9, because the more similar cell types in CNS6 resulted in fewer differentially expressed genes.

### Training

The goal of the training was to generate a collection of profiles of pseudo-bulk cell types used in the composition estimation procedure (Eq. 1). The training was performed in iterations until either the desired performance or the maximum number of iterations was reached. Two sequential training steps were taken in each iteration: (1) on mixed pseudo-bulk samples and (2) on real bulk samples, both with known compositions. A detailed description of the training protocol is provided in the supplementary material. Briefly, top performing sets of profiles in the mixed pseudo-bulk training were further ranked and filtered in the real bulk training. The performance was evaluated by the root mean square error (RMSE) between the expected and predicted proportions:

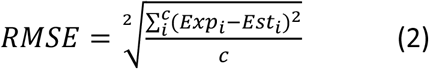

Where *Exp*_*i*_ and *Est*_*i*_ are the expected and estimated proportions of cell type *i* when the evaluation is done across a total number of *c* different cell types of a sample. At the end of the training, multiple sets of profiles were kept for deconvolution.

### Deconvolution

An ensemble method was adopted to integrate the deconvolution results from different sets of profiles to optimize the match between the profiles and the query data. For each query bulk sample, a final cell type composition was calculated from the top *N* estimated compositions based on their gene expression goodness-of-fit RMSEs:

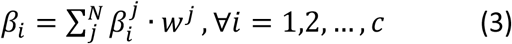

Where 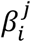 is the proportion of *i*-th cell type in *j*-th estimated composition; the weight *w*^*j*^ was calculated based on the goodness-of-fit *RMSE*_*f*_ of each set of profiles (i.e., *RMSE* from observed and fitted values of gene expression in Equation (1)):

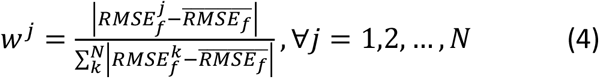

The results presented here were generated with N=5, that is the lowest 5 RMSE values from all profile sets.

### Stepwise deconvolution

Attempting to decompose samples into a very large number of cell types can be difficult as multiple small fractions are hard to predict accurately. To avoid this pitfall, we envisioned a multi-step process: in the first step, the major cell types of interest would be queried, followed by more refined deconvolution steps querying a narrower set of cell types or subtypes based on the outcome of the first step. Following this design, we curated two groups of datasets to include the cell types of interest. The first group consisted of 17 datasets to account for 9 major cell types (Major9), including astrocytes, cardiomyocytes, endothelial cells, hepatocytes, macrophages, neurons, oligodendrocytes, pancreatic and skeletal muscle cells. These are the main cell types of interest for neurodegenerative disease research as well as unintended types that might arise as off-target cell fates. The second group of 9 datasets focused on cell types specific for the central nervous system (CNS). From this dataset, we derived two sets of profiles: the CNS6 that included neurons, astrocytes, endothelial cells, microglia, oligodendrocytes and pericytes, as well as Neuron3 representing neuronal progenitors, inhibitory and excitatory neurons (Fig. 2).

### Comparison to other deconvolution approaches

The mixed pseudo-bulk samples with known cell type proportions that were generated as part of the CellMap pipeline were also used to compare the performance of CellMap to three publicly available methods: MuSiC^13^, SCDC^15^ and Bisque^14^.

Even though multiple sc/sn datasets were used for the profile generation, the input to the CellMap deconvolution step was a reduced data bundle comprised of the cell type profiles. The other three methods (MuSiC, SCDC and Bisque) work directly with sc/sn datasets instead of the pre-generated profiles. To avoid loading all input sc/sn datasets into the memory for the comparisons, an expression matrix was created by merging subsets of cells from them. At least half of the cells were randomly selected for each cell type from each dataset, not to exceed 20M cells of a certain type. Three such input sc/sn expression matrices were created corresponding to the three CellMap profile sets.

The implementations of MuSiC, SCDC and BiSque were slightly modified from their published versions deposited in GitHub. The changes included adjustments to the output format and enabling parallel computing. The modified applications are also available from GitHub together with the expression matrices of the pseudo-bulk samples used for the benchmarking.

The comparison was performed for each cell type separately. The RMSE values were calculated across each cell type by comparing the expected and predicted compositions of the pure and mixed pseudo-bulk training samples. The RMSE was also computed for each pseudo-bulk sample across all cell types of a cell type group (Major9, CNS6, Neuron3).

### Training datasets

The sc/sn RNA-seq datasets (bulk RNA-seq for muscle cells) used to generate the cell type profiles were also used for the pseudo-bulk training (Supplementary Table 1). Multiple rounds of random selections of cells minimized the overlap between the cells used for profile generation (from pseudo-bulk pure samples) and training (on pseudo-bulk mixed samples). Diseased samples were excluded when appropriate annotations were provided. For consistency, we reran the RNA-seq pipeline on these datasets when the raw data was available, as noted in Supplementary Table 1. In addition to these pseudo-bulk samples, true bulk samples from public repositories were used for the true bulk training.

### Testing datasets

To evaluate the performance of CellMap, we assembled a collection of true bulk RNA-seq datasets of purified primary cells and iPSC-derived cell lines, independent from the training sets used in CellMap. For the datasets with primary cells, the information provided by the authors about the cell type composition of the samples was accepted to be the ground truth. In the case of the iPSC-derived cells, the entire target cell type was used as expected composition for the purpose of computing an RMSE. Even though these RMSE values reflected more on the deviation of cell line from the target cell type than on the performance of CellMap, they were deemed to be useful for detecting the changes in composition and similarity of iPSC derived cells relative to the primary cell types. In addition, we used bulk RNA-seq data from brain tissues of the ROSMAP dataset with matching immunohistochemistry (IHC) and snRNA-seq data as ground truth^18, 19^ (Supplementary Table 2).

## Results

### Normalization across datasets

As revealed by principal component analysis (PCA), the pure pseudo bulk samples without any prior normalization tended to cluster by data sources rather than by cell types (Fig. 3A, B, Supplementary Figures S1-S2). This was not surprising, given the differences in sequencing library preparation and various sc/sn sequencing platforms employed to generate the datasets. This strong batch effect would have adversely affected the performance of the decomposition algorithm. Depending on the presence of overlapping cell types among datasets, two different ComBat normalization strategies were deployed to eliminate batch effects. The overlap of cell types among the reference datasets used to generate the Major9 cell type profiles was sparse (Figure 2A). In this scenario, ComBat was applied to the pseudo-bulk pure samples for each cell type separately across their source datasets. The pseudo-bulk samples, that clustered by sequencing platforms originally, became clearly grouped by cell types after this batch correction (Fig. 3D). This strategy proved to be necessary because a normalization across all datasets and all cell types tended to remove cell type specific expression patterns (Fig. 3C). On the contrary, the datasets used for the CN6 and Neuron3 profiles had highly overlapping cell types (Fig. 2B, C). In this case, ComBat normalization across all pseudo-bulk samples in one step was sufficient to remove the batch effects without reducing signals due to real cell type differences. As shown on the corresponding PCA plots, the samples that originally grouped by datasets and sequencing technologies were properly clustered by cell types after normalization (Supplementary Figures S1-S2).

**Figure 3.**
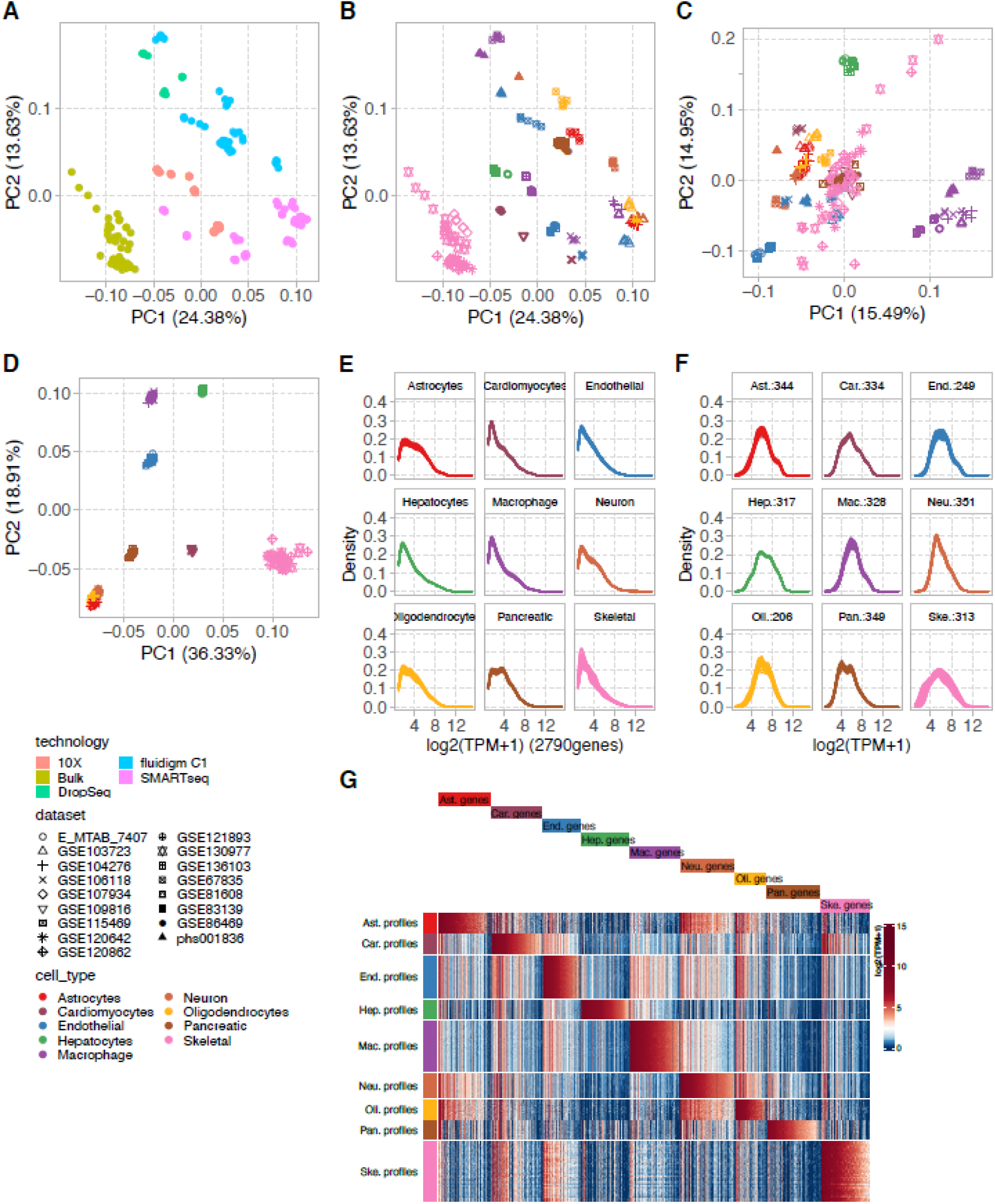
Major9 cell type profiles. (A, B) The first two principal components of the pseudo-bulk expression profiles of the Major9 cell types before normalization colored by sequencing platforms of the source datasets (A) and by cell types (B) showed the samples grouping by sequencing platforms. (C) After ComBat normalization across all datasets, cell type information was lost. (D) After ComBat normalization separately on each cell type, the pseudo-bulk samples grouped by cell types, as expected. (E) Expression profiles of all genes in the pure pseudo-bulk samples. (F) A subset of profile genes was selected such that their expression levels were comparable across cell types. (G) The expression of the profile genes was higher in their respective cell type compared to all other types.

### Performance on the training data

Using the Major9 profiles, CellMap predicted the composition of the pseudo-bulk mixed samples with median RMSE values below 0.1 for all the 9 cell types (Fig. 4A). The lowest performance was observed on endothelial cells and macrophages. When computed across cell types in each dataset, the median RMSE was below 0.1 for the pseudo bulk samples generated from all but one dataset (Fig 4B). The median RMSE was the highest for the true bulk samples (Fig. 4B last column). This latter is not surprising because, even though these true bulk samples contain purified cell types, they are rarely 100% pure and homogeneous.

**Figure 4.**
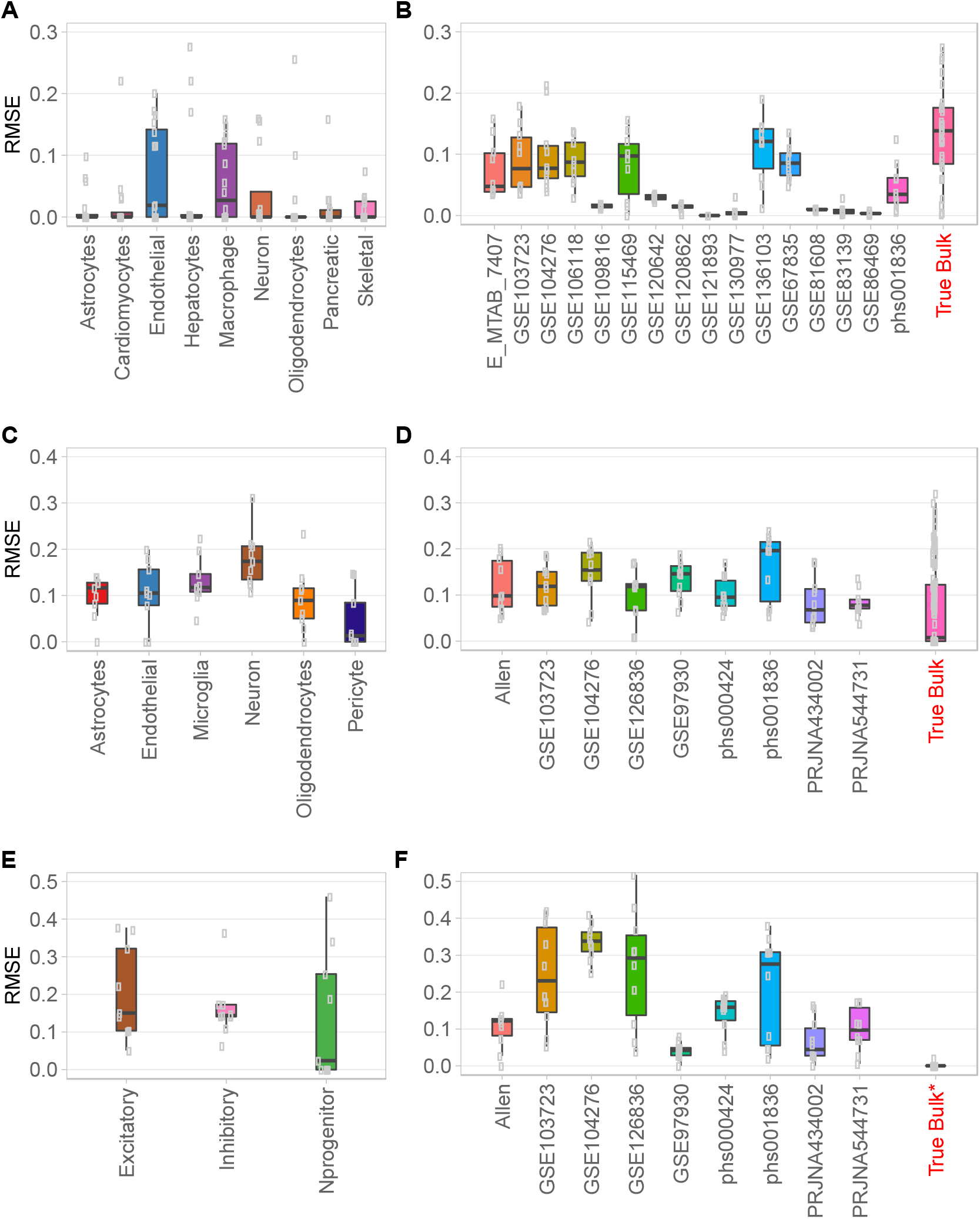
The performance of CellMap on pseudo-bulk mixed samples and true bulk using the three sets of profiles. (A.B) Datasets used for Major9 profiles; (C, D) for CNS6 profiles; (E, F) for Neuron3 profiles. (A, C, E) RMSE values by cell types with each data point representing one dataset; (B, D, F) RMSE values by test datasets with each data point representing one pseudo-bulk mixed or one true bulk sample.

The predictions using the CNS6 profiles were slightly less accurate on average. Having to combine single-cell and single-nuclei datasets for the CNS6 profile set increased the variability of the expression profiles. As a result, predicting the neuronal fractions accurately proved to be more challenging, with RMSE values reaching 0.2 (Fig. 4C). The prediction accuracy across datasets was relatively similar since they tended to have most of the CNS6 cell types: four datasets had median RMSE below 0.1 and five datasets had median RMSE between 0.1 - 0.2. The median RMSE of the predictions of the true bulk samples was well below 0.1, but the individual RMSE values fell into a wide range from 0.01 to slightly over 0.3 (Fig. 4D).

Predicting the neuronal subtypes in Neuron3 proved to be the most difficult task. The median RMSE values ranged from 0.02 to 0.15 for the three subtypes: inhibitory, excitatory and progenitors (Fig. 4E). On a dataset-by-dataset level, 4 datasets yielded accurate predictions with median RMSEs at or below 0.1, while the other 5 had higher RMSE values with larger spread (Fig. 4F). We attribute this increased difficulty to the often-subtle differences between the expression profiles of inhibitory and excitatory neurons that sometimes result in inconsistent labeling of these neuronal subtypes in different datasets. Cell type identification in scRNA-seq dataset clusters, whether it is done manually or by automated annotation tools, is a challenge because the clusters are not completely homogeneous. Slight differences in the selection of cluster marker genes can lead to discrepancies in cell type annotations, especially when these cell types are very similar to each-other, as pointed out by authors working on methods for automated annotation of cell types in sc/sn RNA-seq data^20, 21^. Furthermore, the neuronal progenitors correspond to a continuum of cells at different stages of maturity which makes their characterization difficult. This is reflected in the variability of the prediction accuracy of these cell types across datasets (Fig. 4E).

### Testing results

The performance of CellMap was evaluated on independent bulk RNA-seq datasets by computing the RMSE between the predicted and expected compositions. We performed the tests with both sets of profiles that contained the cell types matching those expected to be found in the test datasets, the Major9 and CNS6. The calculated RMSE values were grouped by purified primary cell types, bulk samples from the ROSMAP dataset, and iPSC-derived cell types. Given a lack of quantitative information about the true cell type content, the expected composition of the iPSC-derived cells was set to 100% of the target cell type in order to allow the calculation of the RMSE.

On the primary cells, the performance of CellMap, as measured by RMSE, was comparable to the performance on the training datasets. Generally, higher RMSE values were obtained on the iPSC-derived cell lines, reflecting their imperfect resemblance to the target primary cell types. It might also be the case that such cell cultures include cells that are not fully differentiated or have stray fates. Median RMSE values were below 0.2 for the purified primary cell type samples using either set of profiles, except for pericytes, while the median RMSE of iPSC-derived samples were below 0.3, except for a subset of iPSC-derived astrocytes (Fig. 5A, B). The neurons and microglia (or macrophages in the case of Major9) were predicted close to the expected 100% purity. The astrocytes and pericytes proved to be the most challenging. The availability of reference purified pericyte datasets was limited. The ones we identified contained less than 100% pericyte-like cells and expressed genes also considered to be fibroblast and oligodendrocyte markers^22^.

**Figure 5.**
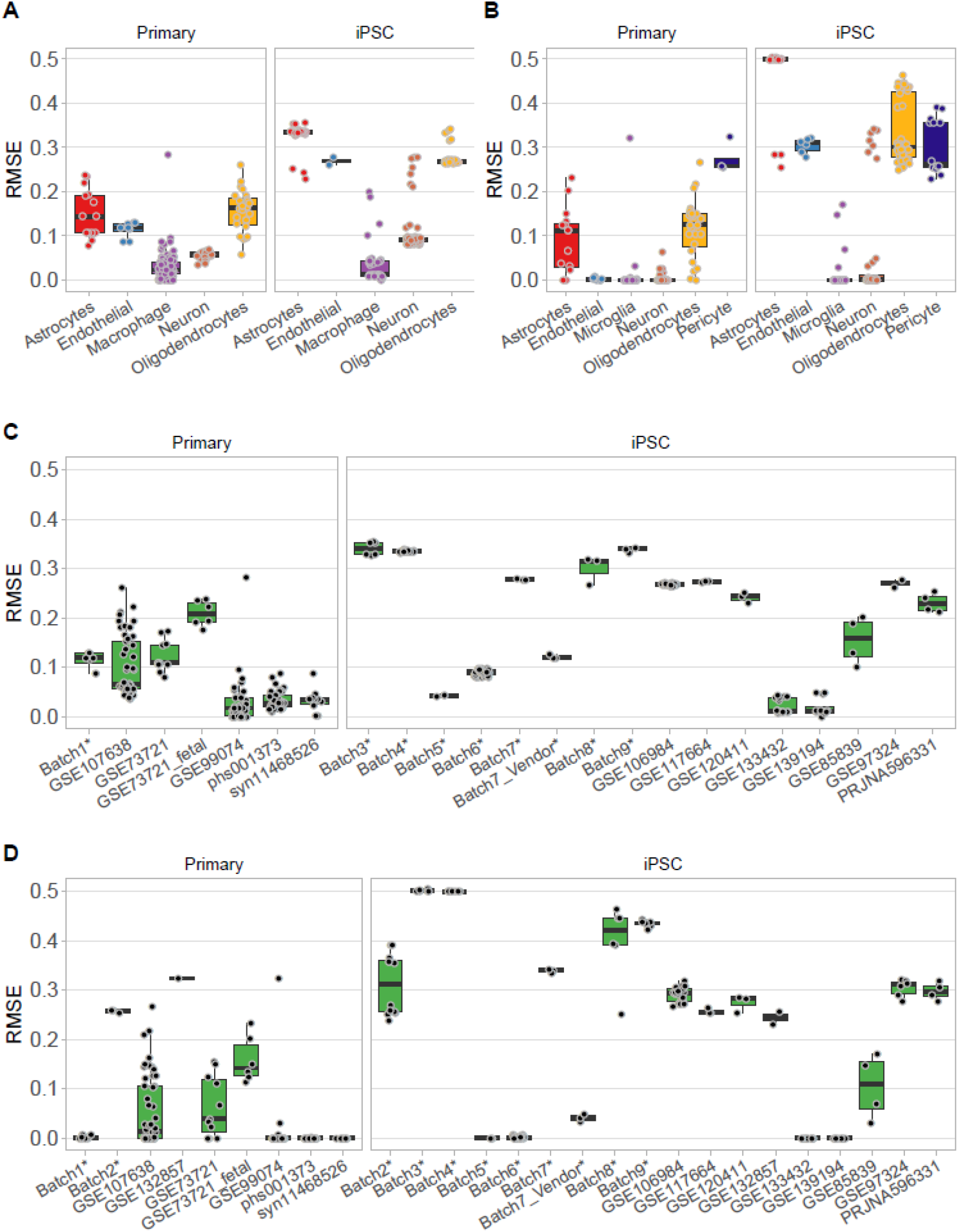
CellMap performance on testing datasets. (A, C) Independent datasets used for the evaluation of the performance by using the Major9 and (B, D) CNS6 cell type profiles. (A, B) RMSE values by cell types with each data point representing one dataset, separated by type of culture into Primary and iPSC datasets. (C, D) RMSE values by datasets with each data point representing one independent reference bulk sample. Datasets denoted by * refer to in-house characterized cell lines (GSE174379).

The differences in performance with the Major9 and CNS6 profile sets confirmed that the input datasets used for generating the cell type expression profiles had a substantial influence on the outcome.

### Examples of CellMap applications

The iPSC-derived cardiomyocyte dataset (GSE122380) was generated from a time course with 16 time points and 19 human cell lines capturing differentiation from iPSCs to mature cardiomyocytes^23^. Despite variable cardiomyocyte purity and marker gene expression levels across samples, CellMap clearly revealed the increasing cardiomyocyte fraction with time (Fig. 6A).

**Figure 6.**
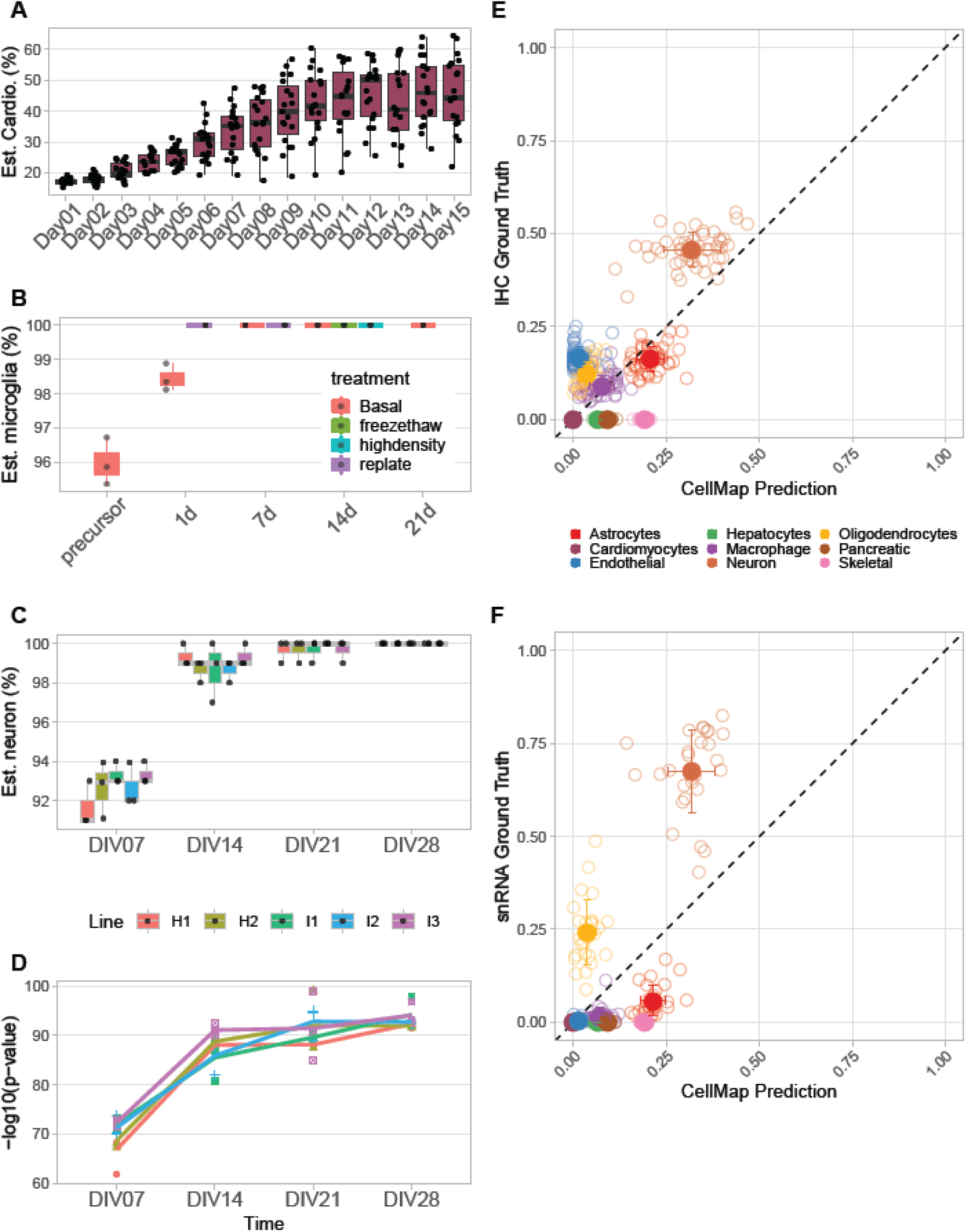
CellMap predictions on bulk RNA-seq data from iPSC-derived cell types and ROSMAP brain samples. (A) Time course of iPSC-derived cardiomyocytes from GSE122380. (B) Multiple treatments of iPSC-derived Microglia. (C, D) Comparing five batches of iPSC derived NGN2 neurons. (E, F) Sample composition predictions for ROSMAP samples relative to composition measured by IHC and snRNA-seq. Error bars represent the standard deviations across samples for each cell type.

Microglia were derived from iPSCs in-house and the effects of different conditions and treatments during differentiation were tested. CellMap was applied to this dataset to determine the optimal time frame for differentiation and to assess how the treatments that such cell lines might have to be subjected to would alter the outcome. CellMap correctly indicated that the precursors were already microglia-like, and the microglia content increased to nearly 100% by day 7 of the differentiation. Additionally, the cells were resistant to both replating and freeze-thaw cycles as indicated by the unaltered microglia composition (Fig. 6B).

Another in-house bulk RNA-seq dataset was generated to assess batch-to-batch variability of iPSC-derived NGN2 neurons similar to those described by Schmid et al. ^24^. The CellMap output indicated low batch-to-batch variability and the cells reaching fully differentiated states by day 21 (Fig. 6C). The results showed not only the increasing predicted percentages of neurons as the cultures differentiated, but also the decreases in p-values, indicating that the maturing cells were acquiring transcriptomic profiles more closely resembling those of the primary neurons.

We applied CellMap to samples from the ROSMAP dataset that had matching immunohistochemistry (IHC) and snRNA-seq data available that we used as the ground truth regarding their cell type compositions^18^. What is accepted as the ground truth also affects the apparent success of the prediction. In the case of the ROSMAP samples, comparing the predicted composition to the IHC data resulted in better predictions than using the cell type information from the matched snRNA-seq data, as there are cell type biases introduced during nuclei isolation (Fig. 6E, F). Similar observations were made previously by Patrick et al.^19^ Based on their analysis and the technical variability of snRNA-seq data, we anticipate the IHC proportions to be closer to the true composition. A good overall correlation was achieved between the predicted cell type fractions and the IHC ground truth. CellMap tended to underpredict the neuronal content, while quite accurately predicted the astrocyte components. The oligodendrocyte and endothelial contents were underestimated, seemingly substituted by other cell types. Endothelial cells are especially easily confounded with other cell types. While they present a set of common features, they also possess considerable heterogeneity depending on their local environment in various organs and tissues^25^. Furthermore, differences in size and RNA content of various cell types can also influence the accuracy of the prediction of cell type proportions^26^.

### Comparison to other deconvolution tools

CellMap performed better than Bisque in each of the categories based on RMSE value, while its performance was very similar or slightly better than that of MuSiC and SCDC in predicting the composition of the pseudo-bulk samples (Fig 7). The range of RMSE values was smallest for CellMap, indicating more consistent predictions across different reference data platforms and cell types. We attribute this robustness to the use of normalization applied to the pseudo-bulk samples as part of the cell type profile generation. In the most difficult task of differentiating between neuronal subtypes (Neuron3 set), CellMap outperformed each of the other three methods (Fig.7C, F). More detailed comparisons by cell types and by input datasets are provided in the supplementary material separated by input datasets used for the Major9, CNS6 and Neuron3 profile sets (Figures S3, S4). While its performance is on par with other existing methods, CellMap has the major advantage of flexibility in using reference datasets with non-overlapping cell types and the ability to expand the cell type repertoire with more cell types of interest as reliable and good quality sn/sc RNA-seq datasets become available.

**Figure 7.**
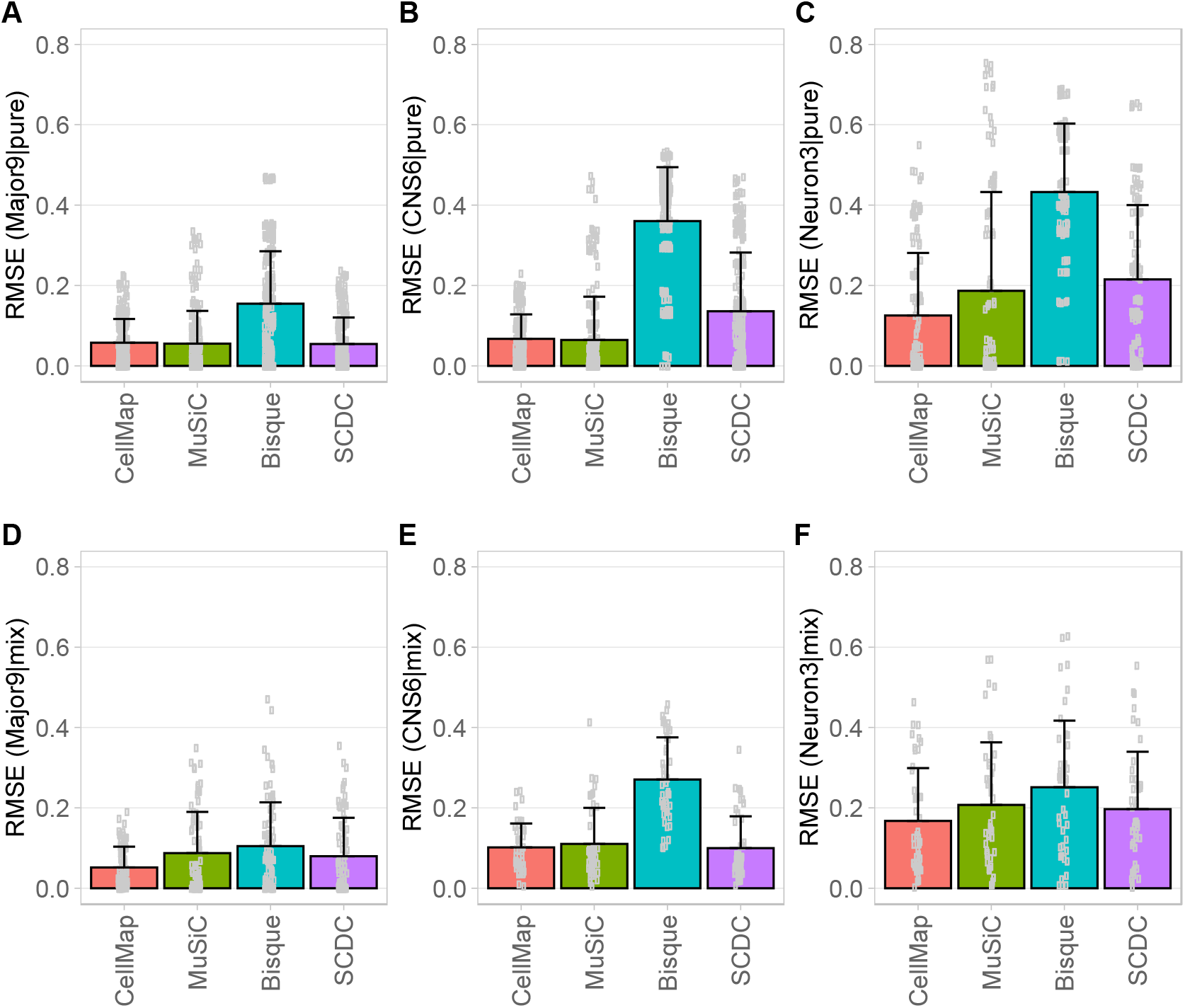
Prediction RMSE values obtained with CellMap, MuSiC, Bisque and SCDC. (A, B, C) Comparisons done using the pseudo-bulk pure samples. (D, E, F) Comparisons across the pseudo-bulk mixed samples. The tests were performed separately on the training datasets used for generating (A, D) the Major9, (B, E) CNS6 and (C, F) Neuron3 profile sets. Each dot represents one pseudo-bulk sample generated with cells from one of the training datasets.

## Discussion

We have demonstrated that CellMap can be applied with good accuracy to evaluate cell composition in bulk RNA-seq data. A key feature of CellMap is its ability to be readily re-trained as new sc/sn RNAseq datasets become available and therefore improve the predictions. This information may come “for free” as CellMap is a downstream analysis on RNA-seq data that may have been generated for other purposes (e.g., differential expression or pathway analysis).

Discrepancies between cell type labeling in different sc/sn datasets can result in inaccurate cell type profiles which have a detrimental effect on the deconvolution accuracy^20, 21^. Instead of discarding such datasets, a possible solution is to test the consistency of cell type assignments across reference datasets with the help of label transfer algorithms and eliminating only the cells that have conflicting annotations^27, 28^.

CellMap was designed with the characterization of human iPSC-derived cell types in mind. However, it is necessary to acknowledge that iPSC-derived cells may not match primary cells exactly. Even primary cells kept in culture for a relative short time differ from their freshly isolated counterparts. When we compared the transcriptome profile of our in-house generated iPSC-derived microglia to the primary microglia dataset generated by Gosselin and his colleagues, it was not surprising to find out that our iPSC-derived cells were more similar to the cultured primary cells than the freshly isolated ones^29^. It would therefore be unreasonable to expect perfect predictions on iPSC-derived cells when training on the transcriptomic profiles of fully differentiated, mature primary cells.

Despite these shortcomings, deconvolution tools like CellMap are very useful and readily deployed in a cell line development workflow. In combination with microscopic observations of morphological features, they allow us to evaluate the progression of the differentiation in a time-course experiment to ensure correct cell fate. They are excellent quality control methods for verifying batch-to-batch consistency of iPSCs and provide valuable guidance in differentiation protocol optimization.

## Data Availability and Computer Code

The CellMap R package, including the R code to generate the manuscript figures, is available from https://github.com/interactivereport/CellMap. The in-house RNA-seq data generated from iPSC-derived cell lines has been deposited in the Gene Expression Omnibus (GEO tracking number GSE174379). All other datasets are publicly available and are listed in the supplementary material.

## Acknowledgements

The authors thank Robin Kleiman and Chao Sun from Biogen for their support and expert advice in cell type characterization and in-house NGS data generation.

## Author contributions

This study was conceived of and led by MIZ, SE, EL and BZ. ZO designed the algorithm, wrote the computer code and performed the data analysis with contributions from NB, XZ and KL. PC and SE lead the production iPSCs and provided scientific insight into the biology of neuronal cell types. PFC and RC performed the RNA-seq data generation. MIZ and ZO wrote the paper with feedback from all coauthors.

## Supplementary Data

### Supplementary Figures

**Figure S1. CN6 cell type profiles**. (A, B) The first two principal components of the pseudo-bulk expression profiles of the CNS6 cell types before normalization colored by sequencing platforms of the source datasets and by cell types showed the samples grouping by sequencing platforms. (C) After cell type ComBat normalization, the pseudo-bulk samples grouped by cell types, as expected. (D) Expression profiles of all genes in the pseudo-bulk samples. (E) A subset of profile genes was selected such that their expression levels were comparable across cell types. (F) The expression of the profile genes was higher in their respective cell type compared to all other types.

**Figure S2. Neuron3 cell type profiles**. (A, B) The first two principal components of the pseudo-bulk expression profiles of the Neuron3 cell types before normalization colored by sequencing platforms of the source datasets and by cell types showed the samples grouping by sequencing platforms. (C) After cell type ComBat normalization, the pseudo-bulk samples grouped by cell types, as expected. (D) Expression profiles of all genes in the pseudo-bulk samples. (E) A subset of profile genes was selected such that their expression levels were comparable across cell types. (F) The expression of the profile genes was higher in their respective cell type compared to all other types.

**Figure S3. Comparison between CellMap, MuSiC, Bisque and SCDC by Major9 cell types**. (A, B, C, D) RMSE values between predicted and expected compositions on the pseudo-bulk pure samples with single cell types. (E, F, G, H) Prediction accuracies measured by RMSE on pseudo-bulk mixed samples. The tests were performed on the training datasets used for generating the Major9 profiles. Each data point represents a dataset.

**Figure S4. Comparison between CellMap, MuSiC, Bisque and SCDC by CNS6 cell types**. (A, B, C, D) RMSE values between predicted and expected compositions on the pseudo-bulk pure samples with single cell types. (E, F, G, H) Prediction accuracies measured by RMSE on pseudo-bulk mixed samples. The tests were performed on the training datasets used for generating the CNS6 profiles. Each data point represents a dataset.

**Figure S5. Comparison between CellMap, MuSiC, Bisque and SCDC by Neuron3 cell types**. (A, B, C, D) RMSE values between predicted and expected compositions on the pseudo-bulk pure samples with single cell types. (E, F, G, H) Prediction accuracies measured by RMSE on pseudo-bulk mixed samples. The tests were performed on the training datasets used for generating the Neuron3 profiles. Each data point represents a dataset.

### Supplementary Tables

**Table 1.** Datasets used as input for cell type profile generation and performance comparison of CellMap, MuSiC, Bisque and SCDC.

**Table 2.** Datasets used for testing.

