## Supplementary Methods and Figures for "CellMap: Characterizing the types and composition of iPSC-derived cells from RNA-seq data"

### Supplementary Material

### Methods

#### *Default settings for generating the pseudo-bulk profiles*

For each cell type, 5 pure pseudo-bulk samples were generated from each input reference dataset. From the same datasets, 10 mixed cell type pseudo-bulk samples were generated. At least 2 million reads were required for each pseudo-bulk. To be included in the expression matrix, a gene had to have an expression level of at least 4 TPM in at least 80% of samples of one of the cell types. The maximum number of training iterations was set to 10. These parameters along with the following ones can be adjusted by the user according to the properties of the datasets used as input for the profile generation and training.

#### *Pseudo-bulk training*

For each of the 10 mixed pseudo-bulk samples derived from one of the sc/sn datasets, 1000 randomly generated combinations of *c* pseudo-bulk pure profiles, ($P_1$, $P_2$, …, $P_c$), selected across all datasets, were used to estimate the cell type proportions of the mixed pseudo-bulk. Since we were creating mixed pseudo-bulks from a given sc/sn dataset, we knew exactly the cell type proportions in the mixture. The root mean square error (RMSE, eq. 2) between the expected and predicted proportions was used to evaluate and rank each of the 1000 candidate profile sets for each mixed pseudo-bulk sample. The top 50 sets of profiles achieving the best predictions for each pseudo-bulk mixture were selected and retained, resulting in a maximum total number of 500 optimal sets of profiles per input dataset, where duplicates might exist due to the random selection picking the same profiles more than one time. Since we carry out this procedure per dataset, in the end we have a maximum of (500 x number of datasets) candidate sets of pseudo-bulk pure profiles.

#### *True bulk training*

When true bulk samples with known cell type composition were available for training, the selected sets from the pseudo-bulk training above were further filtered based on their performance on these true bulk samples. For each of the bulk training samples (i.e., true bulk samples with known composition), a final RMSE (eq. 2) was calculated between the expected and the weighted estimated (eq. 3) cell type compositions. If it was higher than 0.1, we filtered out a number of candidate profile sets that had poor individual RMSEs. The poorer the final composition RMSE was for a bulk training sample, the more of its lower ranking profile sets were removed. The number was determined by its final composition RMSE compared to others:

$$N_k^{rm} = \frac{RMSE_k}{\sum_k RMSE_k} \cdot N^{rm}, \forall k\{k|RMSE_k > 0.1\}$$

where $N_k^{rm}$ is the number of lower ranking profile sets to be removed for the *k*-th bulk samples; $RMSE_k$ is the final RMSE for the *k*-th bulk sample; $N^{rm}$ is the maximum number of profile sets to be removed. We choose 75% of total selected profile sets from pseudo training to be $N^{rm}$, the upper limit for removal.

If true bulk samples with known cell type compositions are not available for this second stage of filtering, we recommend the user to pick a smaller subset, less than the top 50 sets, from the pseudo-bulk training procedure (previous section) or assemble pseudo-bulk from profile-independent sc/sn RNAseq dataset with high quality.

#### *Training iterations*

The profile generation, normalization and training (pseudo bulk and real bulk) were repeated until either the final RMSE for all bulk samples was smaller than 0.1 or the maximum iteration (10) was reached. On the second and subsequent iterations, the profile sets generated from the prior iteration were appended to the current iteration's pseudo bulk mixture training profile sets and the combined set was sent into the real bulk training procedure (see Figure 1.).

However, if every real bulk which included a cell type achieved an RMSE smaller than 0.1, this cell type would not be included in the pseudo bulk mixtures for next iteration training, as they no longer needed further optimization. In this way, if we had cell types that were sufficiently well estimated, we no longer created pseudo bulk training samples for them.

### Supplementary Figures

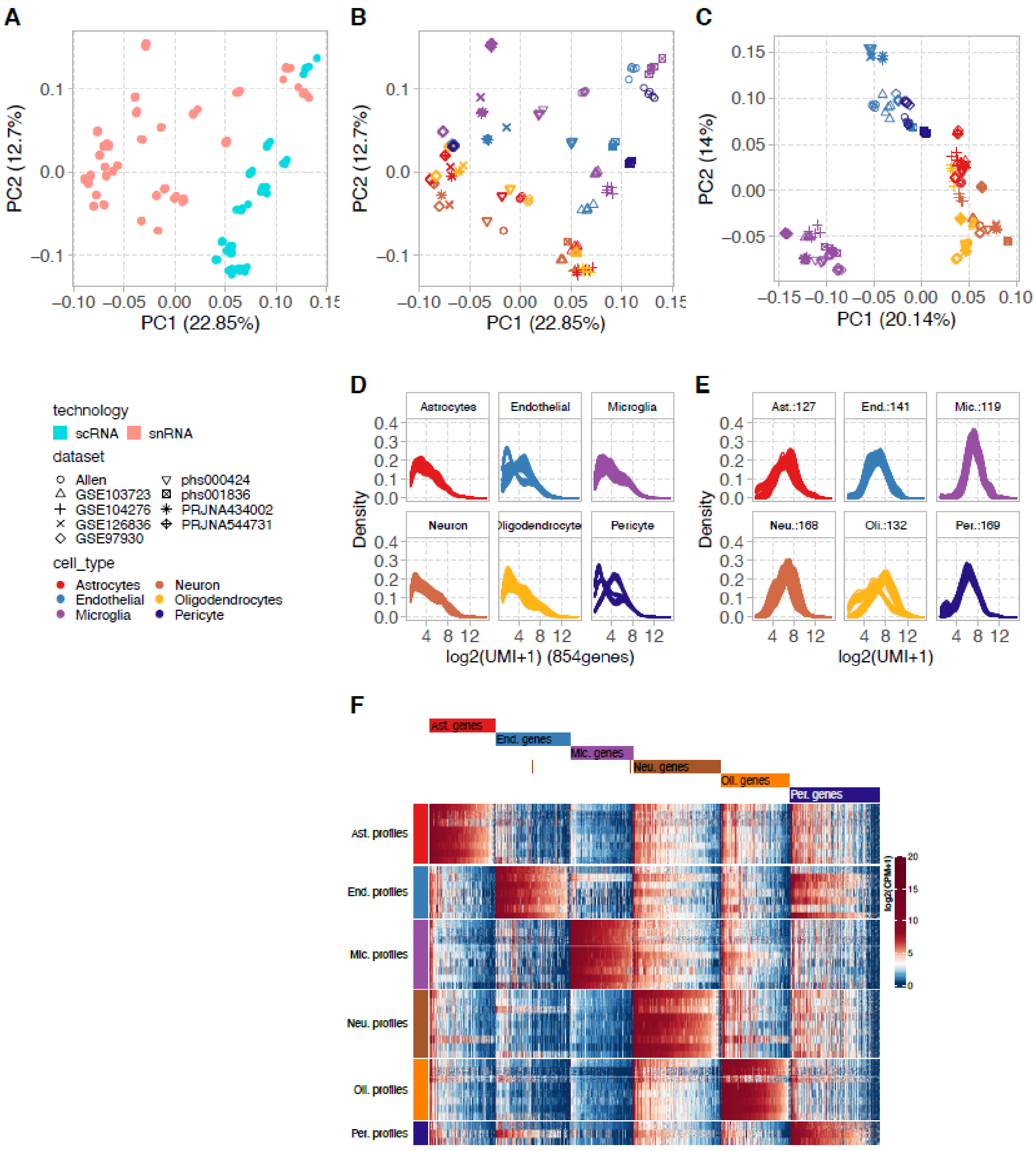


**Figure S1. CN6 cell type profiles.** (A, B) The first two principal components of the pseudo-bulk expression profiles of the CNS6 cell types before normalization colored by sequencing platforms of the source datasets and by cell types showed the samples grouping by sequencing platforms. (C) After cell type ComBat normalization, the pseudo-bulk samples grouped by cell types, as expected. (D) Expression profiles of all genes in the pseudo-bulk samples. (E) A subset of profile genes was selected such that their expression levels were comparable across cell types. (F) The expression of the profile genes was higher in their respective cell type compared to all other types.

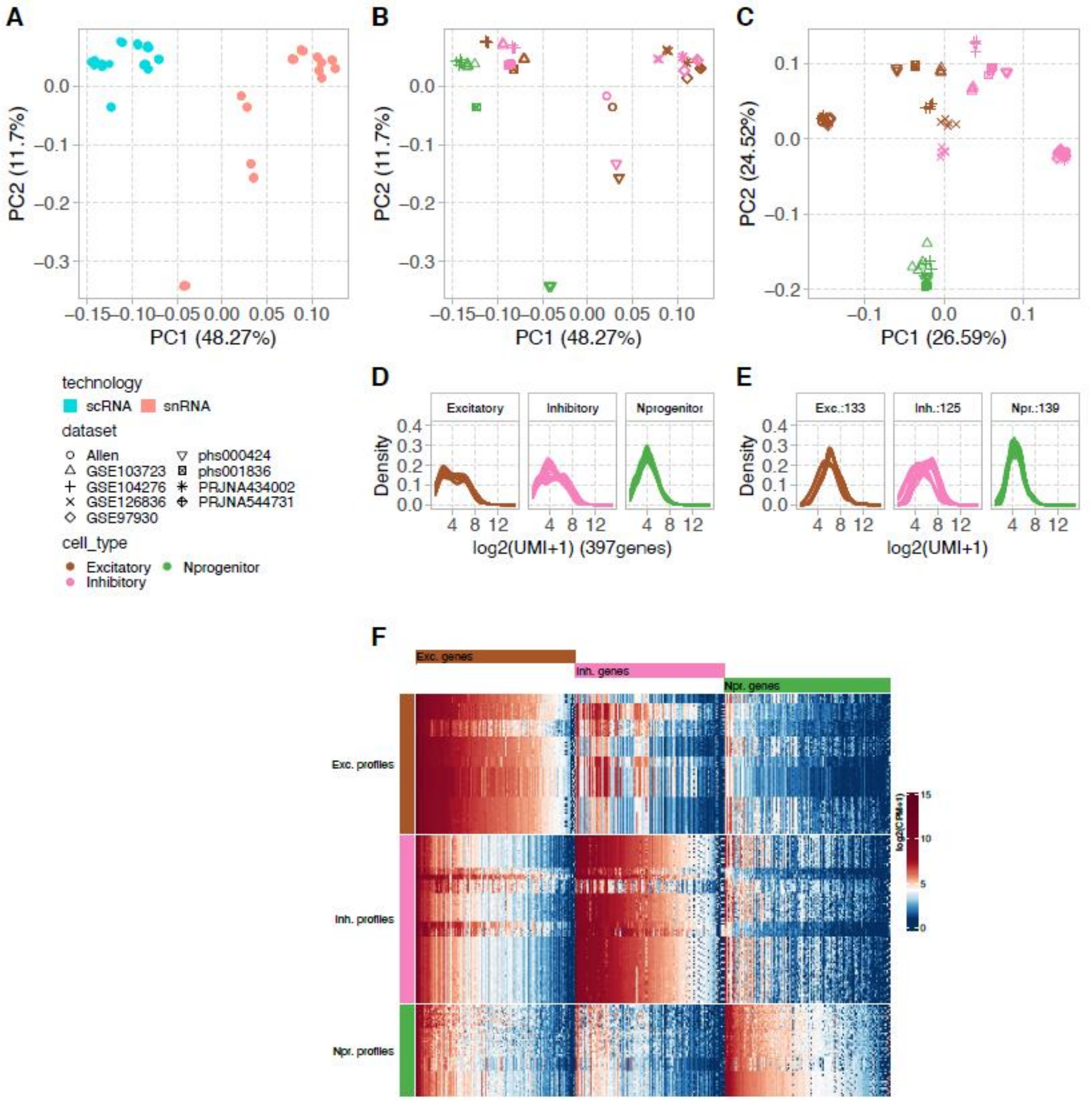


**Figure S2. Neuron3 cell type profiles.** (A, B) The first two principal components of the pseudo-bulk expression profiles of the Neuron3 cell types before normalization colored by sequencing platforms of the source datasets and by cell types showed the samples grouping by sequencing platforms. (C) After cell type ComBat normalization, the pseudo-bulk samples grouped by cell types, as expected. (D) Expression profiles of all genes in the pseudo-bulk samples. (E) A subset of profile genes was selected such that their expression levels were comparable across cell types. (F) The expression of the profile genes was higher in their respective cell type compared to all other types.

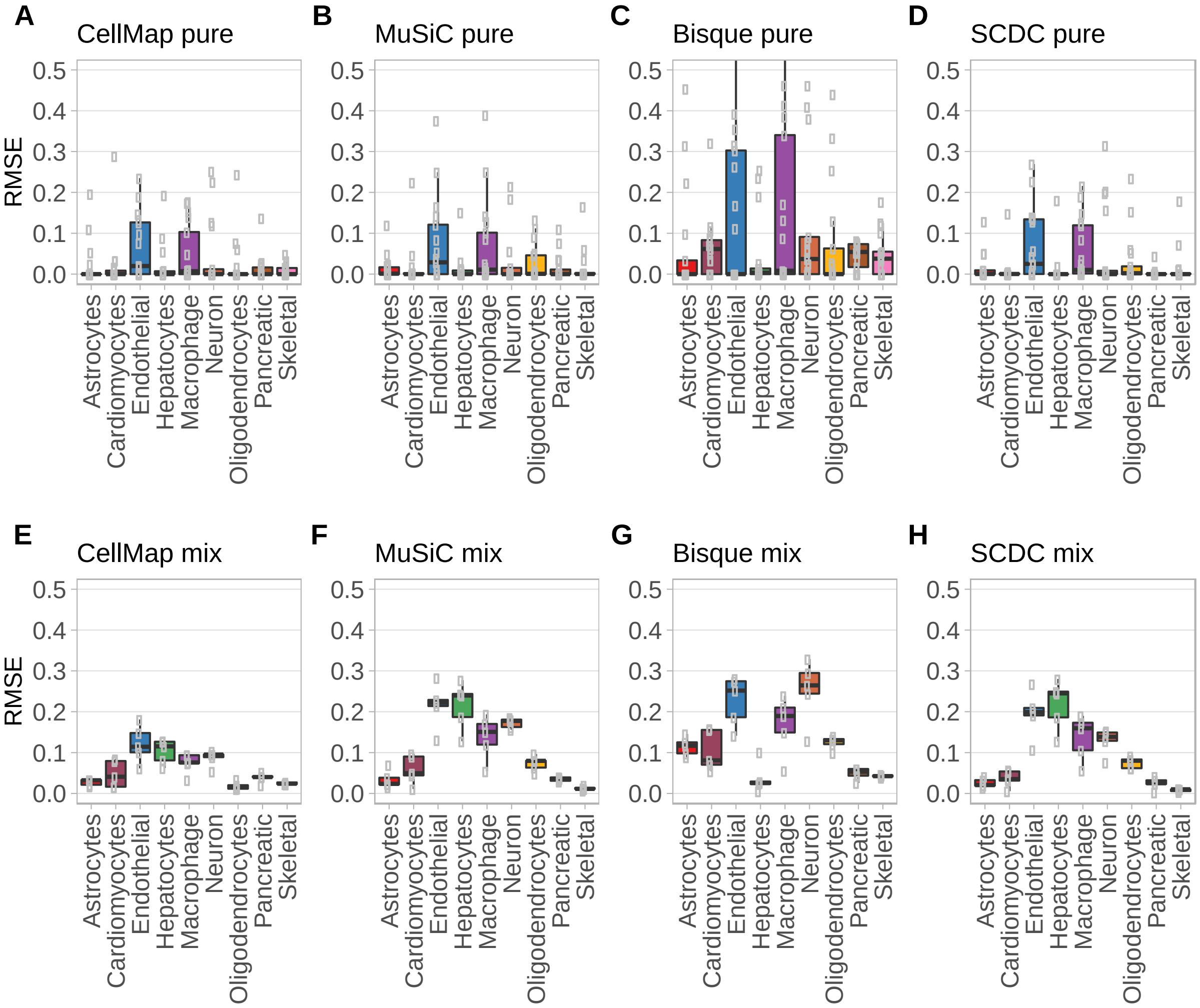


**Figure S3. Comparison between CellMap, MuSiC, Bisque and SCDC by Major9 cell types**. (A, B, C, D) RMSE values between predicted and expected compositions on the pseudo-bulk pure samples with single cell types. (E, F, G, H) Prediction accuracies measured by RMSE on pseudo-bulk mixed samples. The tests were performed on the training datasets used for generating the Major9 profiles. Each data point represents a dataset.

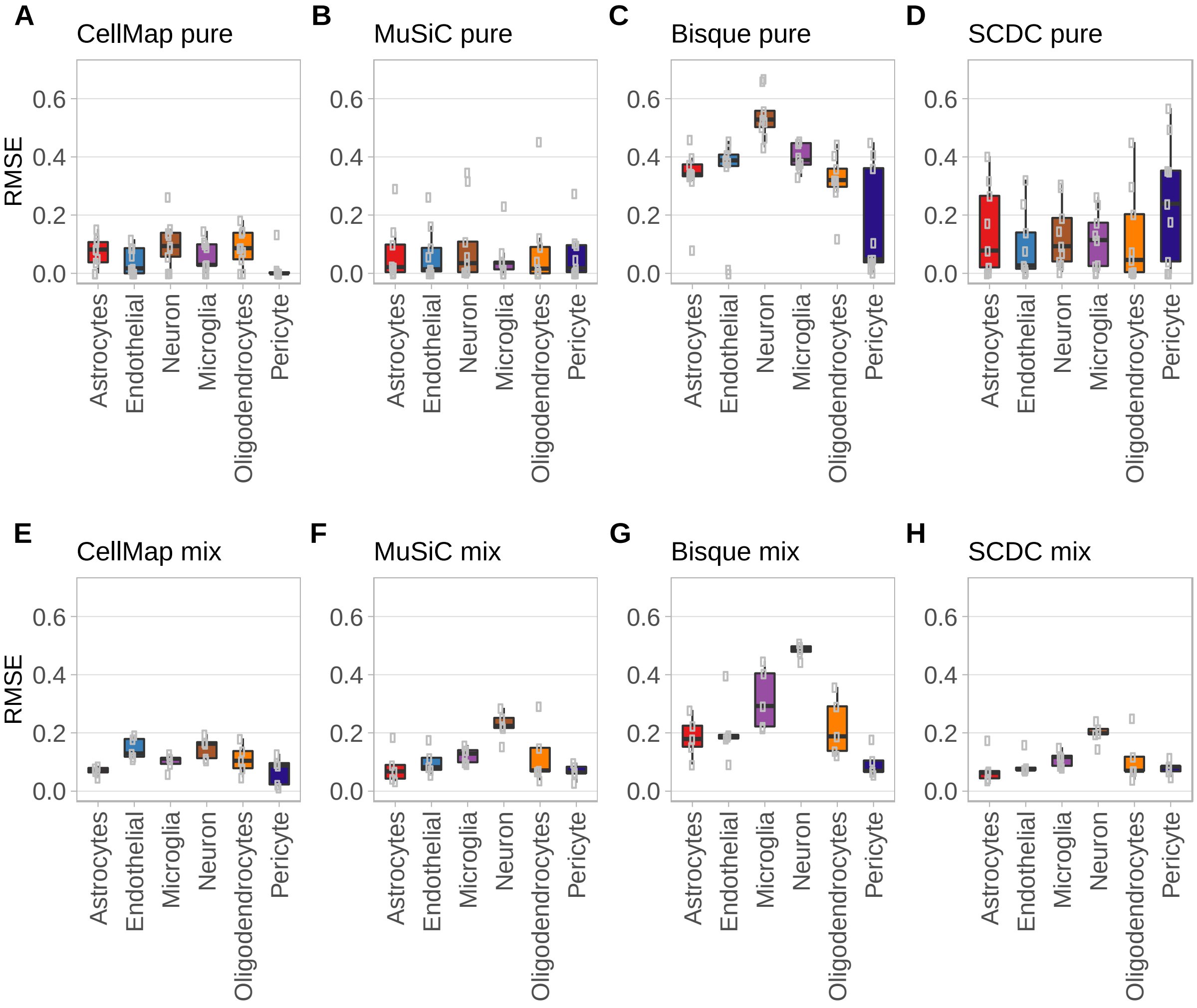


**Figure S4. Comparison between CellMap, MuSiC, Bisque and SCDC by CNS6 cell types.** (A, B, C, D) RMSE values between predicted and expected compositions on the pseudo-bulk pure samples with single cell types. (E, F, G, H) Prediction accuracies measured by RMSE on pseudo-bulk mixed samples. The tests were performed on the training datasets used for generating the CNS6 profiles. Each data point represents a dataset.

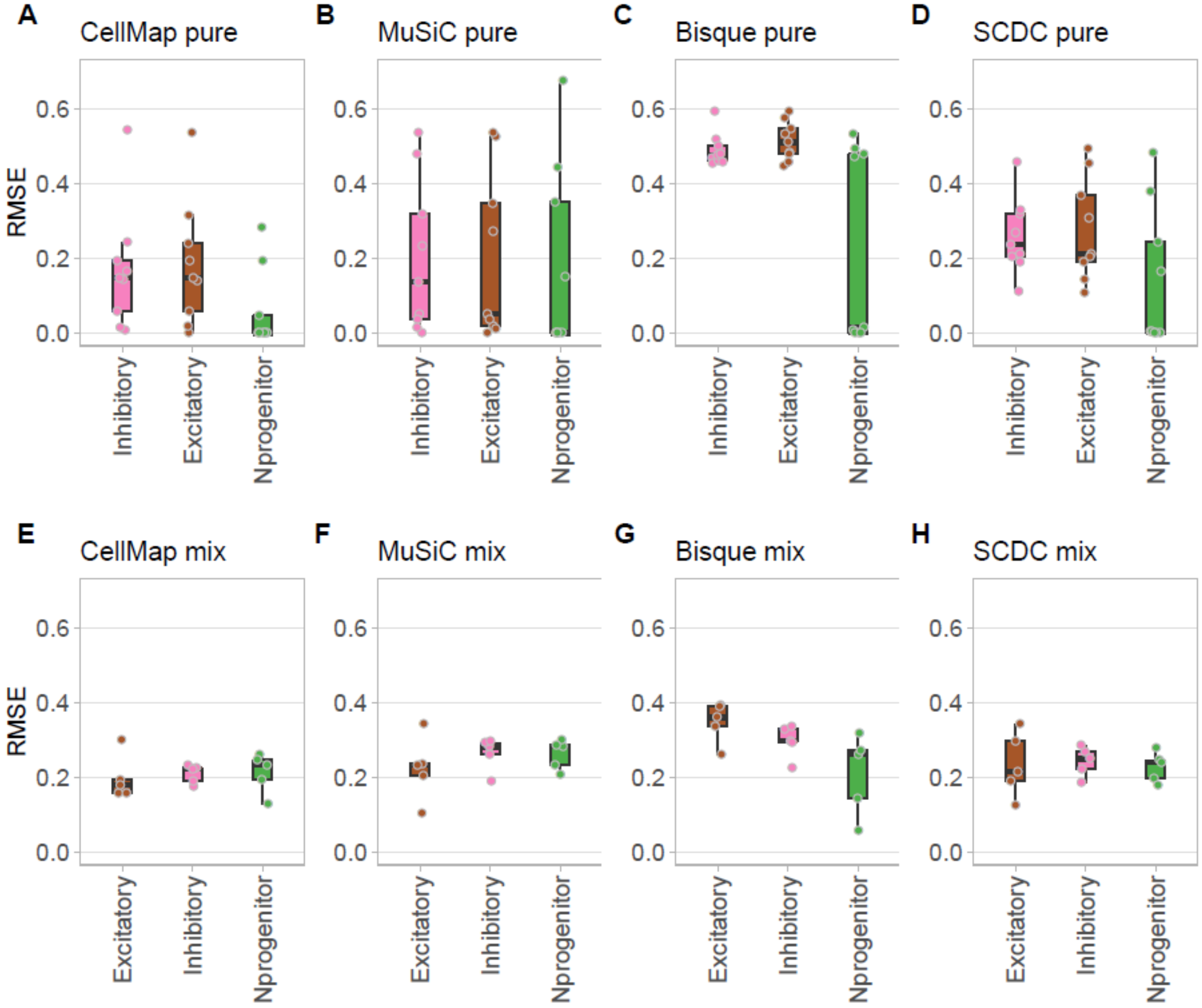


**Figure S5. Comparison between CellMap, MuSiC, Bisque and SCDC by Neuron3 cell types.** (A, B, C, D) RMSE values between predicted and expected compositions on the pseudo-bulk pure samples with single cell types. (E, F, G, H) Prediction accuracies measured by RMSE on pseudo-bulk mixed samples. The tests were performed on the training datasets used for generating the Neuron3 profiles. Each data point represents a dataset.
